# An Atlas of Immune Cell Exhaustion in HIV-Infected Individuals Revealed by Single-Cell Transcriptomics

**DOI:** 10.1101/678763

**Authors:** Shaobo Wang, Qiong Zhang, Hui Hui, Kriti Agrawal, Maile Ann Young Karris, Tariq M. Rana

## Abstract

Chronic infection with human immunodeficiency virus (HIV) can cause progressive loss of immune cell function, or exhaustion, which impairs control of virus replication. However, little is known about the development and maintenance, as well as heterogeneity of immune cell exhaustion. Here, we investigated the effects of HIV infection on immune cell exhaustion at the transcriptomic level by analyzing single-cell RNA sequencing of peripheral blood mononuclear cells from two healthy subjects (15,121 cells) and six HIV-infected donors (28,610 cells). We identified nine immune cell clusters and eight T cell subclusters according to their unique gene expression programs; three of these (exhausted CD4^+^ and CD8^+^ T cells and interferon-responsive CD8^+^ T cells) were detected only in samples from HIV-infected donors. An inhibitory receptor KLRG1 was identified in the exhausted T cell populations and further characterized in HIV infected individuals. We identified a novel HIV-1 specific exhausted CD8^+^ T cell population expressing KLRG1, TIGIT, and T-bet^dim^Eomes^hi^ markers. *Ex-vivo* antibody blockade of KLRG1 restored the function of HIV-specific exhausted CD8^+^ T cells demonstrating the contribution of KLRG1^+^ population to T cell exhaustion and providing a novel target for developing immunotherapy to treat HIV chronic infection. Analysis of gene signatures also revealed impairment of B cell and NK cell function in HIV-infected donors. These data provide a comprehensive analysis of gene signatures associated with immune cell exhaustion during HIV infection, which could be useful in understanding exhaustion mechanisms and developing new cure therapies.

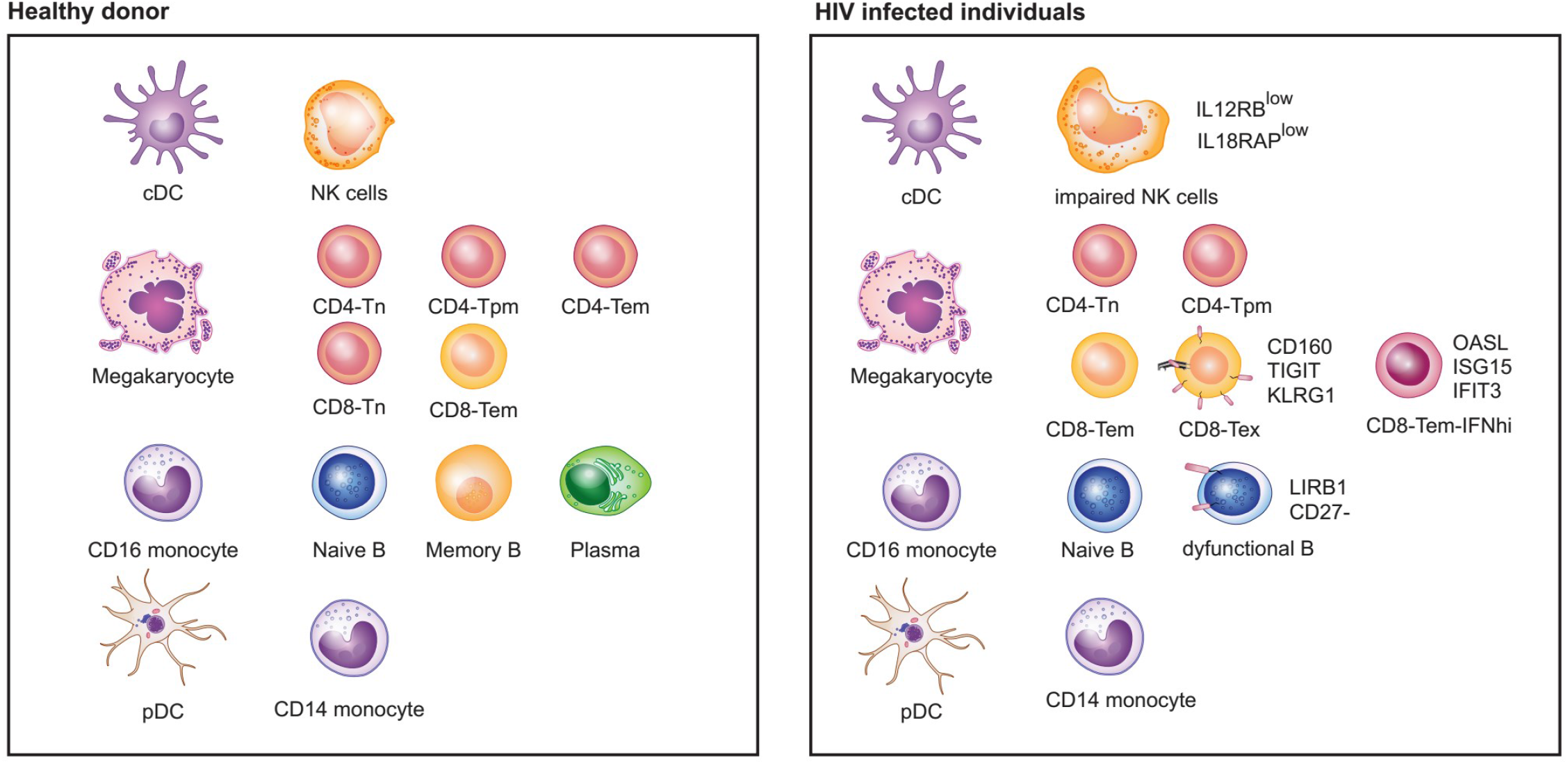

## INTRODUCTION

More than 76 million people have been infected with human immunodeficiency virus (HIV) since the epidemic was first recognized in the 1980s. Approximately 37 million people worldwide are currently living with HIV infection, of whom only ~21 million have access to antiretroviral therapy (UNAIDS.org; http://www.unaids.org/en/resources/fact-sheet). The development of potent combination antiretroviral therapy (cART) has allowed HIV viremia to be controlled and significantly reduced the mortality of HIV-infected individuals; however, withdrawal of treatment leads to a rapid rebound of viremia, indicating that the host immune system remains unable to control viral replication ^1,2^. HIV-induced dysfunction of the host immune system is one of the major causes of this recrudescence. Persistent exposure to viral antigens leads to chronic activation of immune cells, progressive loss of function, and ultimately, to a state of exhaustion accompanied by complete loss of effector function ^3–6^. For example, exhausted CD8^+^ T cells lose their cytotoxic effector function and the ability to eradicate HIV-infected cells ^7^. In addition, CD8^+^ T cell exhaustion prevents the differentiation of effector cells into memory cells with the ability to undergo rapid reactivation upon encounter with antigen ^7,8^. Chronically activated CD4^+^ T cells also lose their effector functions, including the production of cytokines, such as IL-2 and IL-21, that sustain HIV-specific CD8^+^ T cells ^9–11^. Consequently, HIV-induced CD4^+^ T cell depletion and exhaustion also results in dysfunction of HIV-specific CD8^+^ T cells, resulting in disease progression ^7,8^. Therefore, understanding the mechanisms by which HIV infection leads to immune exhaustion is critical for the development of vaccines and therapies for HIV and, possibly, for other viral infections.

Sustained expression of high levels of inhibitory receptors such as PD-1, CTLA-4, CD160, TIGIT, and TIM-3 is a hallmark of T cell exhaustion. In HIV-infected individuals, expression of multiple inhibitory receptors has been demonstrated to correlate positively with plasma viral load and disease progression ^6,12–15^. Notably, neutralizing antibody-mediated blockade of these receptors reverses T cell exhaustion by increasing the production of effector molecules and the proliferation of HIV-specific CD4^+^ and CD8^+^ T cells ^6,12–15^. Blockade of PD-1 in simian immunodeficiency virus (SIV)-infected macaques results in expansion of SIV-specific CD8^+^ T cells, proliferation of memory B cells, decreased viremia, and prolonged survival ^16,17^. Similarly, treatment with a PD-L1-blocking antibody suppresses viremia and increases CD4^+^ T cell counts in chronically HIV-infected humanized mice ^18^. Intriguingly, a recent report described that nivolumab (anti-PD-1) treatment of an HIV-infected patient with non-small-cell lung cancer restored the function of HIV-specific CD8^+^ T cells and decreased the HIV reservoir ^19^. A clinical trial is currently ongoing to evaluate the safety and efficacy of PD-1 blockade in HIV-infected individuals with a low CD4^+^ T cell count (NCT03367754).

Despite the great promise of reversal of T cell exhaustion as a curative strategy for HIV, little is known about the mechanisms involved in immune cell exhaustion in HIV-infected individuals at the transcriptomic level. In addition, chronic HIV infection is associated with exhaustion not only of T cells but also of B and NK cells ^20–22^. Therefore, it is possible that these three exhausted cell populations could cooperate in contributing to host immune system dysfunction, allowing unchecked HIV replication and disease progression.

Single-cell RNA sequencing (scRNA-seq) is a powerful technological advance to analyze the transcriptomic profiles of individual cells, thus enabling the heterogeneity of cells affecting biological processes to be analyzed ^23^. scRNA-seq can reveal cellular identity as well as spatial organization and clonal distribution in complex heterogeneous immune populations ^24,25^. The complexity of the immune system is a major obstacle to understanding the host immune response to HIV infection. The diversity and dynamic states of individual immune cells cannot be precisely revealed by traditional bulk expression profiles, which provide averaged values from highly heterogeneous populations ^26^. In addition, there are limitations to identifying, isolating, and analyzing special or new cell populations, especially for the rare cell subtypes ^27^. scRNA-seq has successfully identified new types of human blood dendritic cells (DCs), monocytes, and progenitors, resulting in a revised taxonomy of blood cells (Villani et al., 2017). Inspired by these studies, we reasoned that scRNA-seq could make it possible to analyze the effects of HIV infection on the landscape of immune cell types, including rare cell populations.

Recently, scRNA-seq technology has been applied to investigate viral diversity, heterogeneity of infection states, latency and reactivation, and virus–host interactions ^28–33^. For example, Cohn et al. identified unique signature genes in reactivated latent CD4^+^ T cells, isolated from HIV infected individuals, which allow cell division without trigging the cell death pathways induced by HIV-1 replication ^28^. In another model of HIV latency, scRNA-seq identified transcriptional heterogeneity in latent and reactivated CD4^+^ cells ^29^. Other scRNA studies provided new insights into transcriptional dynamics in Zika and dengue viral infections ^33^ and during cytomegalovirus latency ^32^, and the heterogeneity of influenza virus infection ^31^. These reports represent proof-of-concept that scRNA technology can provide unique insights into the virus–host interaction.

In this study, we performed scRNA-seq on PBMCs from HIV-infected and healthy donors to determine how HIV infection shapes the landscape of exhausted immune cells. We found that HIV infection induced the appearance of several cell clusters with novel gene signatures, including subsets of exhausted CD8^+^ and CD4^+^ T cells and CD8^+^ T cells that are highly responsive to interferon (IFN). One of the exhausted T cell signature genes was the inhibitory receptor KLRG1, which was coexpressed with TIGIT in exhausted T cells. Blockade of KLRG1 efficiently restored HIV-specific immune responses providing a promising novel target to develop immunotherapy for HIV infection. Finally, we performed integrated analysis of PBMCs and showed that HIV infection also induces B and NK cell dysfunction, which, together with T cell exhaustion, may contribute to HIV-related immunodeficiency.

## RESULTS and DISCUSSION

### Atlas of PBMCs in Healthy and HIV-Infected Donors

To examine the landscape of immune cell exhaustion induced by HIV infection, we isolated PBMCs from healthy and HIV-infected donors and performed scRNA-seq to identify cell clusters by expression cell type-specific gene signatures. Previous studies have suggested that expression of exhaustion genes such as *PD-1, CTLA-4, CD160, TIGIT*, and *TIM-3* is associated with high HIV loads ^12,13^. Therefore, we included three donors with a high viral load and three with a low viral load (>100,000 and <1000 RNA copies/ml of plasma, respectively) in the analysis. A total of 12,852 and 15,758 PBMCs were sequenced from donors with high and low viral loads (hereafter referred to as HL-HIV-infected and LL-HIV-infected donors), respectively (4000-5500 PBMCs/donor). For comparison, we performed scRNA-seq of PBMCs from one healthy donor (ID HD_1) and obtained scRNA-seq data from another healthy donor (ID HD_2; 10X Genomics), to give a final healthy donor dataset from 15,121 PBMCs. An overview of the approach is given in Figure 1A. Through unbiased analysis, we identified nine major cell clusters present in the healthy and HIV-infected donors based on expression of a unique gene signature: CD4^+^ T cells (*CD3D^+^ CD8A^−^ IL7R^hi^*), CD8^+^ T cells (*CD3D^+^ CD8A^+^*), natural killer cells (NK; *CD3D^−^ CD8A^−^ IL7R^−^ GNLY^hi^*), B cells (*MS4A1^+^*), CD14^+^ monocytes (*LYZ^hi^ CD14^hi^*), CD16^+^ monocytes (*LYZ^hi^ FCGR3A^hi^*), conventional dendritic cells (cDCs; *LYZ^hi^ FCER1A^hi^*), plasmacytoid dendritic cells (pDCs; *LYZ^low^ IGJ^hi^*), and megakaryocytes (Mk; *PPBP*^+^). This information is displayed as t-Distributed Stochastic Neighbor Embedding (tSNE) plots of the cell clusters in Figures 1B–D and S1B-F, and expression of the gene signatures defining each cluster is shown as heatmaps and violin plots in Figure S1A. The absolute numbers and percentages of cell subsets in the PBMC samples for each donor are given in Figure S1G. The healthy donor PBMC samples contained the expected proportions of major white blood cell classes: ~50% T cells (CD4^+^:CD8^+^ T cells ~2:1), 10–15% B cells, ~10% NK cells, ~20% monocytes, and ~3% DCs (Figure S1G).

**Figure 1.**
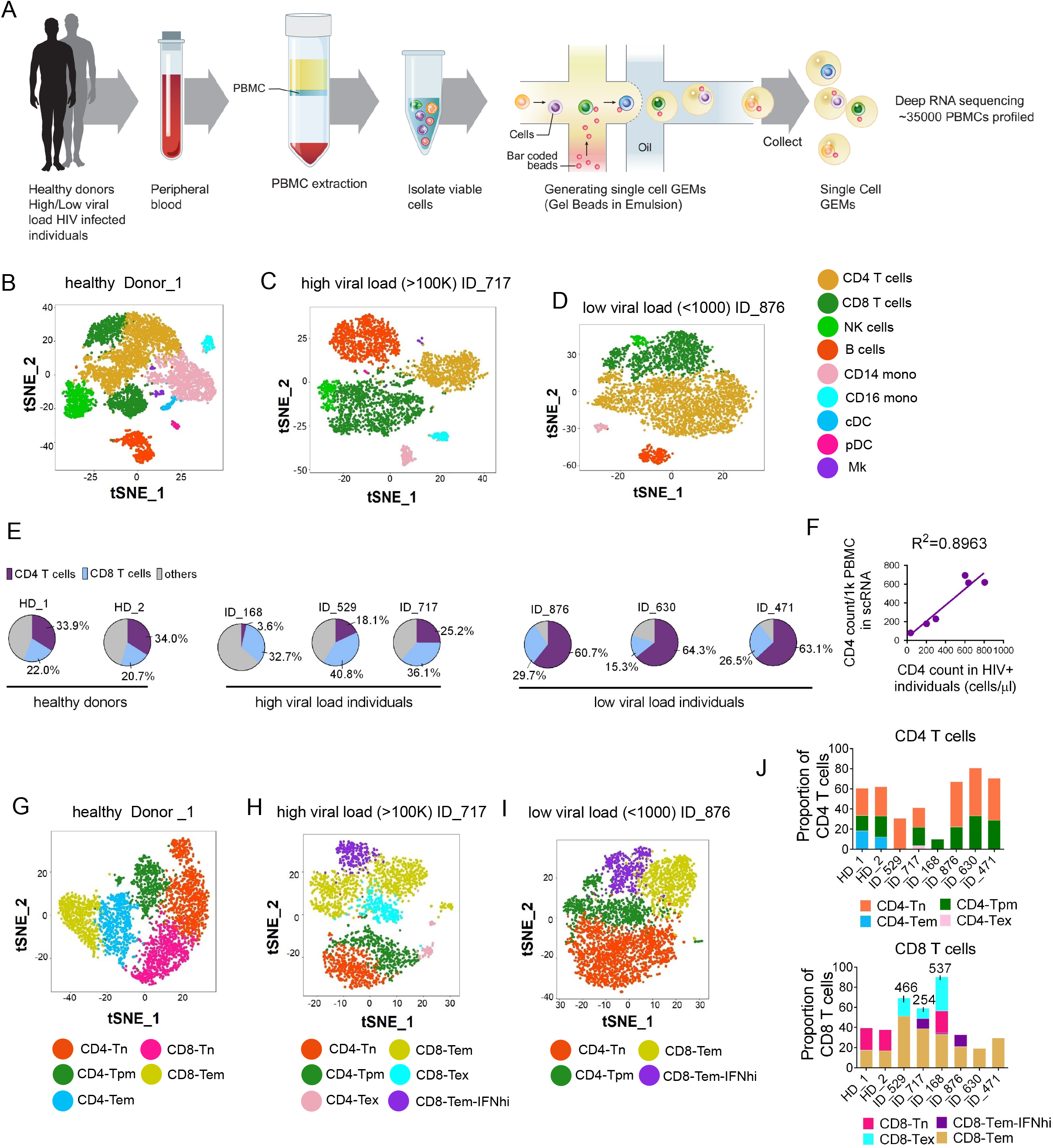
Distinct Cell Clusters can be identified by scRNA-seq of PBMCs from Healthy and HIV-Infected Donors. (A) Overview of workflow. PBMCs were isolated from healthy donors and HIV-infected donors (three each with high and low viral loads [>100,000 and <1000 RNA copies/ml plasma, respectively]). Single cells were captured by gel beads with primers and barcoded oligonucleotides and subjected to deep RNA-seq. (B–D) t-Distributed Stochastic Neighbor Embedding (t-SNE) projection of PBMCs from healthy donor HD_1 (B), high-load HIV-infected donor ID_717 (C), and low-load HIV-infected donor ID_876 (D), showing major cell clusters based on normalized expression of cell type-specific markers. NK, natural killer cells; CD14 mono, CD14^+^ monocytes; CD16 mono: CD16+ monocytes; cDC, conventional dendritic cell; pDC, plasmacytoid dendritic cell; Mk, megakaryocytes. (E) Pie charts showing the percentage CD4^+^ T cells, CD8^+^ T cells, and other PBMC subsets in the healthy and HIV-infected donors indicated in (B–D). (F) Linear regression analysis showing the correlation between CD4^+^ T cell counts calculated from scRNA analysis (cells/1000 PBMCs) vs flow cytometry (cells/μl) of PBMCs from HIV-infected donors. (G–I) tSNE projections for T cell subsets from healthy donor HD_1 (G), HL-HIV-infected donor ID_717 (H), and LL-HIV-infected donor ID_876 (I). Tn, naïve; Tpm, precursor memory; Tem, effector memory; Tex, exhausted; IFNhi, highly IFN-responsive. (J) Percentage of the indicated subclusters of CD4^+^ and CD8^+^ T cells from two healthy donor samples (HD_1, _2), three HL-HIV-infected donors (ID_529, _717, and _168), and LL-HIV-infected donors (ID_876, _630, and _471). See also Figure S1 and S2

Consistent with the known effects of HIV infection, the absolute number and percentage CD4^+^ T cell counts were considerably lower in the three HL-HIV-infected donors (18.1%, 25.2%, 3.6%) compared with the two healthy donors (33.9%, 34.0%). In concert, the percentage (but not absolute cell numbers) of CD8^+^ T cells was increased in the HL-HIV-infected donors (40.8%, 36.1%, 32.7% versus 22.0%, 20.6%; Figures 1E and S1G). We also observed a striking increase in the percentage CD4^+^ T cells in the LL-HIV-infected donor samples (60.7%, 64.3%, 63.1%) (Figures 1E and S1G). Importantly, the absolute CD4^+^ T cell counts in the clinical blood samples and the numbers estimated from the scRNA-seq analysis showed a strong correlation (R^2^=0.8963, Figure 1F), indicating that the scRNA-seq datasets accurately reflect the cell clusters present in the original blood samples.

To identify which T cell subsets are selectively depleted or expanded in the HIV-infected donors, we performed unbiased clustering based on subset-specific marker genes. In the healthy donor PBMCs, three CD4^+^ T cell and two CD8^+^ T cell clusters (Figures 1G, S2A, and S2C) were identified based on the relative enrichment or depletion of signature genes as naïve CD4^+^ T cells (CD4-Tn: *CD8A^−^ CCR7+ IL7R^hi^)*, effector memory CD4^+^ T cells (CD4-T_em_: *CD8A^−^ IL7R^hi^ CCR7^−^ GZMA*^+^) ^34^, and a cluster that we defined as precursor memory cells (CD4-Tpm: *CD8A^−^ IL7R^hi^ CCR7^low^ LTB^hi^*). The putative CD4-Tpm cluster showed a similar signature to that of CD4-Tn cells (e.g., *TCF7, FOXP1*) but they did not express effector function-associated genes (e.g., *GZMA, CCR5, NKG7*), suggesting that they may represent a transitional state between naïve and effector memory status (Gattinoni et al., 2011; Youngblood et al., 2017). This cluster also harbored high expression of the TNF family molecule lymphotoxin β (*LTB*; Figure S2A), which is consistent with a recently described CD4-Tpm cytotoxic cell cluster ^35^. In addition to the three CD4^+^ T clusters, PBMCs from the healthy donors also contained clusters consistent with naïve CD8^+^ T cells (CD8-Tn: *CD8A^+^ CCR7^hi^)* and effector memory CD8^+^ T cells (CD8-Tem: *CD8A^+^IL7R^−^ CCR7^−^ GZMA^+^ NKG7^+^*; Figures 1G, S2A, and S2C).

Notably, the composition and proportion of T cell subtypes in PBMCs were markedly altered by HIV infection. Samples from the donors with high viral loads contained dramatically smaller populations of CD4-Tem and CD8-Tn, and three new cell clusters with unique gene signatures could be discerned; namely, exhausted memory CD8^+^ T cells (CD8-Tex), exhausted memory CD4^+^ T cells (CD4-Tex), and a population of CD8^+^ Tem cells with marked upregulation of IFN-response genes, which we termed CD8-Tem-IFN^hi^ (Figures 1H, S2B, S2D, and S2E). The Tex subpopulations were classified based on previous work showing that CD8^+^ T cells expressing high levels of the exhaustion markers *CD160* and *TIGIT* were enriched in HIV-infected individuals ^14,15^. In the three high-load HIV-infected donors, we found that 18.1%, 10.1%, and 33.9% of total CD8^+^ T cells carried the Tex gene signature. Similarly, CD4-Tex cells were characterized by expression of the exhaustion markers *TIGIT* and *CTLA4*. This is consistent with an earlier demonstration that *CTLA4* is enriched in CD4-Tex cells during HIV chronic infection ^13^. Finally, cells within the CD8-Tem-IFN^hi^ cluster showed enrichment of IFN-stimulated genes such as *OASL, ISG15, IFIT2, and IFIT3*, consistent with expansion of the host antiviral immune response (Figures 1H, 1I, S2E). Interestingly, analysis of samples from the LL-HIV-infected donors showed a reduction in the CD4-Tem and CD8-Tn clusters and the appearance of a CD8-Tem-IFN^hi^ cluster, similar to the observations in samples from HL-HIV-infected donors; however, there were no obvious CD4^+^ or CD8^+^ Tex cell populations in samples from these donors (Figures 1I, 1J, S2E, and S2F).

Collectively, these data identify the major PBMC and T cell subsets affected by HIV infection, including CD4^+^ and CD8^+^ Tex populations and highly IFN-responsive CD8^+^ Tem cells, each of which have unique signature profiles.

### Identification of Novel Genes Associated with T Cell Exhaustion

Since little is known about the gene signatures of Tex cells in HIV-infected donors, we further characterized the CD8^+^ Tex subsets by comparing their gene expression patterns with those of their unexhausted counterparts, which are typically Tem cells (Figure 2A and Table 1). From this analysis, we identified a total of 39 genes that were commonly altered in CD8-Tex cells from at least two of the three HL-HIV-infected donors (p < 0.01, roc test; Figures 2B and 2C). The 24 up-regulated genes in HL-HIV-infected donors included the known exhaustion markers *TIGIT* and *CD160* (Figure 2D) ^14,15^. These inhibitory receptors have also been identified as T cell exhaustion markers in pathogen-infected and tumor-bearing animals ^14,15,36–40^. CD8-Tex cells from the HL-HIV-infected donors also showed up-regulated expression of another inhibitory receptor, killer cell lectin-like receptor subfamily G member 1 (KLRG1), which contains an immunoreceptor tyrosine-based inhibitory motif in the cytoplasmic domain (Figure 2D). KLRG1 is known to be a T cell differentiation marker ^41^. Other genes specifically up-regulated in CD8-Tex cells from HL-HIV-infected donors included *CMC1, SH2D1A, COMMD6, TRAPPC1*, and *COTL-1* (Figures 2B and 2E). Conversely, there were 15 genes commonly down-regulated in CD8-Tex cells from HL-HIV-infected donors, including *IRF1, ITGB1*, and genes encoding the cytotoxic T cell effector molecules granzyme B (*GZMB*), perforin 1 (*PRF1*), and granulysin (*GNLY*) (Figures 2C and 2E). These observations strongly suggest that CD8-Tex cells are less cytotoxic than normal CD8^+^ Tem cells.

**Figure 2.**
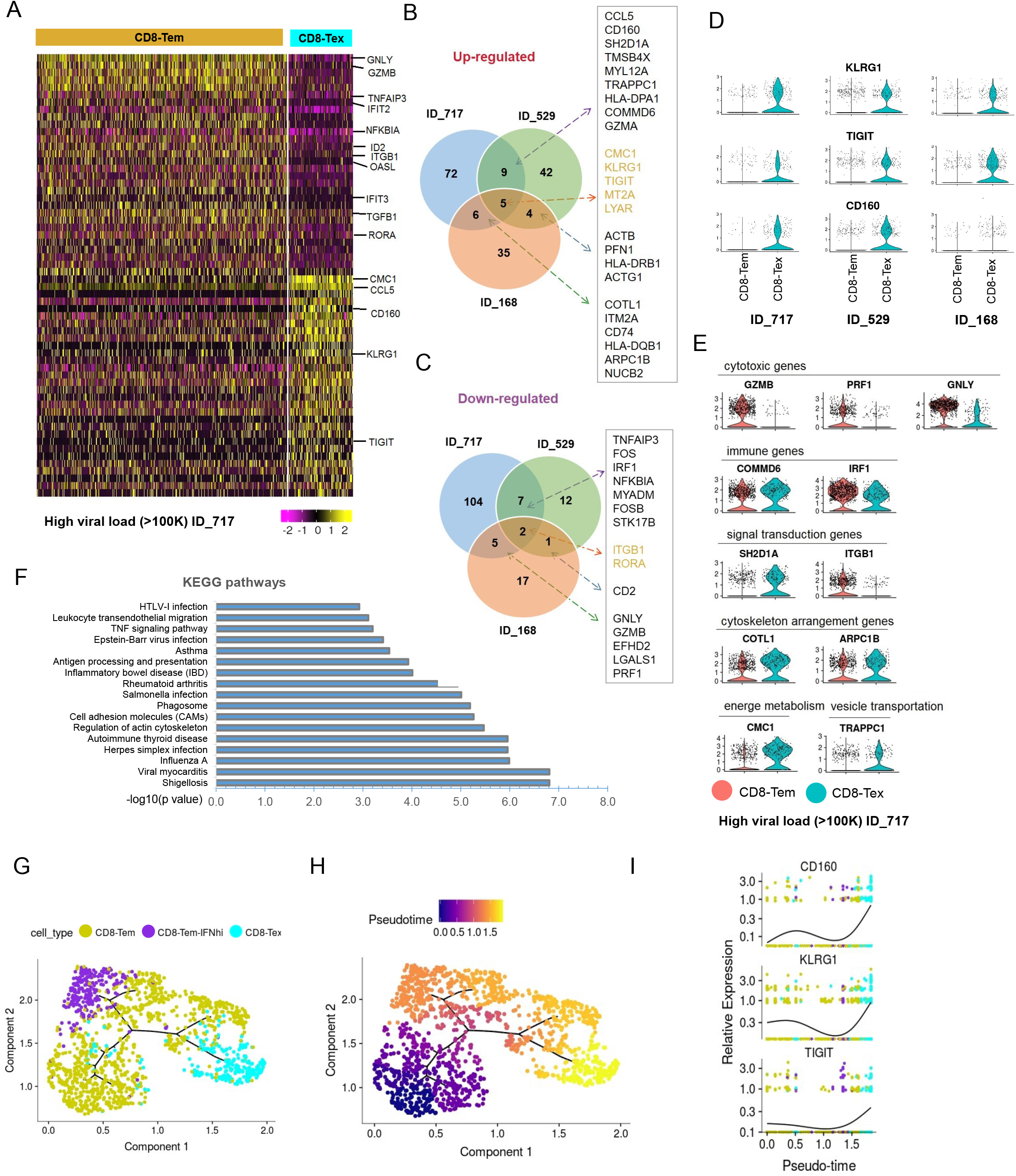
Identification of Novel Signature Genes in CD8-Tex Cells from HIV-Infected Donors. (A) Heatmap showing differentially expressed genes in CD8-Tem and CD8-Tex cells from HL-HIV-infected donor ID_717. The signature genes are indicated to the right of the heatmap. The color code below the map indicates the relative expression levels. (B and C) Venn graphs of conserved up-regulated (B) and down-regulated (C) genes in CD8-Tex cells from the three HL-HIV-infected donors. (D) Violin plots of the conserved up-regulated exhaustion-associated signature genes in CD8-Tex compared with CD8-Tem cells from the three HL-HIV-infected donors. Each dot represents a single cell and the shapes represent the expression distribution. The remaining cells are depicted on the x axis. (E) Violin plots of the up-regulated and down-regulated genes associated with the indicated functions in CD8-Tex compared with CD8-Tem cells from HIV-infected donor (ID_717). Each dot represents a single cell and the shapes represent the expression distribution. The remaining cells are depicted on the x axis. (F) GO analysis of the common genes in exhausted CD8^+^ memory T cells. (G and H) The trajectory plots of the pseudotime analysis shows two distinct trajectories of the CD8-Tem cells in HIV infected individuals. The start of the pseudotime was set to be CD8-Tem cells. The trajectory plots were visualized and colored by cell types (G) and pseudotime (H) separately. (I) Three representative genes CD160, KLRG1, and TIGIT were identified by pseudotime analysis to be significantly enriched in the CD8-Tex cells.

Other immune response genes differentially expressed in HL-HIV-infected donor-derived Tex cells compared with Tem cells included *COMMD6*, an NF-κB-inhibiting protein, and the IFN-responsive transcription factor *IRF1*, which were up-regulated and down-regulated, respectively, in CD8-Tex cells, consistent with HIV-induced suppression of immune signaling (Figure 2E) ^42^. In support of this, *SH2D1A*, which encodes the signaling inhibitor SLAM-associated protein (SAP), was also up-regulated in Tex compared with Tem cells ^43^. SAP has important roles in signaling for T cell differentiation and the antiviral immune response ^43^. Tex cells exhibited reduced expression of the integrin β1-encoding gene *ITGB1*, which has many crucial immune functions and mediates cell–cell interactions ^44^, suggesting another mechanism that may contribute to the exhaustion phenotype. Interestingly, Tex cells showed altered expression of genes involved in cytoskeletal function (*TMSB4X, PFN1, ACTG1, COTL1, ARPC1B*), energy metabolism (*CMC1*), and vesicle transportation (*TRAPPC1*) (Figure 2E). These pathways have also been associated with CD8^+^ T cell exhaustion during lymphocytic choriomeningitis virus infection ^40^. Gene Ontology analysis revealed that the differentially expressed genes common to Tex cells in HIV-infected donors were enriched in pathways involved in pathogen infection, actin cytoskeleton regulation, cell adhesion, antigen processing and presentation, and TNF signaling (Figure 2F).

The development of normal CD8 Tem cell into exhausted T cell is was analyzed by “pseudotime” analysis as the cells differentiate asynchronously and heterogeneously. CD8 effector memory cells were ordered based on single cell transcriptomics by pseudotime analysis using monocle 3 ^45,46^. Root state is defined as normal effector memory. CD8 effector memory cells branched into interferon high effector memory and exhausted cells, revealing a bifurcating trajectory of CD8 differentiation in HIV infected individuals. Exhausted cells are at the end of pseudotime, suggesting a terminal differentiation state (Figures 2G and H). Importantly, this exhaustion sub-branch was occupied by cells expressing high levels of exhaustion genes, CD160, KLRG1, and TIGIT. (Figure 2I). Taken together, these results identified novel signatures of *KLRG1, CD160*, and *TIGIT* characterizes exhausted CD8^+^ T cells in HIV-infected donors.

### KLRG1 blockade effectively restores the function of HIV-specific CD8^+^ T cells

The observation that *KLRG1* was strongly up-regulated and co-expressed with two known HIV induced exhaustion markers *TIGIT, CD160* in Tex cell clusters from all three HL-HIV-infected donors raised the possibility that *KLRG1* may be involved in T cell exhaustion. To test this hypothesis, we analyzed KLRG1 expression in PBMCs from healthy and HIV-infected donors by flow cytometry (Figure 3A and B). We found the frequency of KLRG1 and TIGIT expressing CD8^+^ T cells significantly increased in PBMCs from HIV-infected individuals (Low VL, 10.8%; High VL, 18.1%) compared with healthy donors (6.6%). Since PBMCs from higher viral load PBMCs showed severe exhaustion, we did see higher ratio of KLRG1^+^TIGIT^+^ populations in high viral load PBMCs. KLRG1 is widely used as a terminal differentiation marker in lymphocytes ^47^. It is necessary to clarify the exact exhausted KLRG1 + subpopulation induced by HIV chronic infection. T-bet and Eomes had been proved to be highly associated with exhausted phenotype induced by HIV infection in a reciprocal pattern ^48^. T-bet^dim^ Eomes^hi^ bulk or HIV-specific CD8^+^ T cells showed poor functional characteristics ^48^. Therefore, we further analyzed the percentages of T-bet^dim^ Eomes^hi^ population in KLRG1^+^TIGIT^+^ CD8^+^ T cells. The result showed that the frequency was much higher in HIV-infected PBMCs (Low VL, 23.3%; High VL, 23.3%) compared with that from healthy donors (HD, 6.0%) (Figure 3C). In addition, in both high viral load and low viral load group, the percentage of T-bet^dim^ Eomes^hi^ cells was much higher in KLRG1^+^TIGIT+ population (potential exhausted) compared with KLRG1^−^TIGIT^−^ (non-exhausted) population (Figure 3C). This data suggests that KLRG1^+^TIGIT^+^ T-bet^dim^ Eomes^hi^ population in CD8^+^ T cells represent a novel exhausted T cell population in HIV chronic infection.

**Figure 3.**
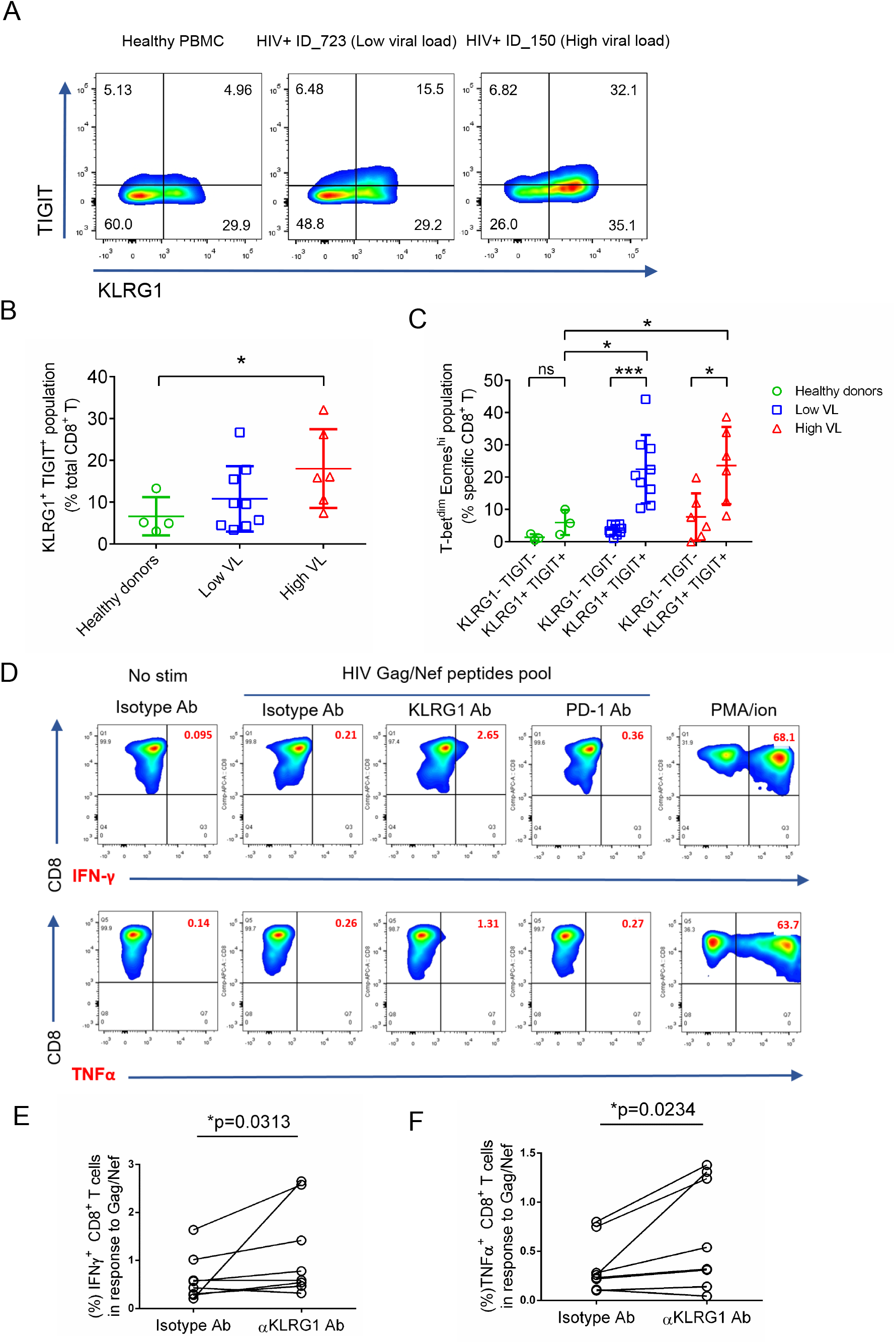
KLRG1 blockade effectively restores the function of HIV-specific CD8^+^ T cells. (A) Flow cytometry of KLRG1^−^ and TIGIT^−^ expressing CD8^+^ T cells from the indicated healthy and HIV-infected donors. Numbers indicate the percentage of KLRG1^+^ TIGIT+, KLRG1^−^ TIGIT^−^, KLRG1^+^ TIGIT^−^ and KLRG1^−^ TIGIT^+^, -expressing cells. (B) KLRG1^+^ TIGIT^+^ population is increased in HIV HL-individuals. The percentage of KLRG1^+^ TIGIT^+^ cells was analysis by flow in healthy donors (n=4), HIV LL- (n=9) and HL- (n=6) individuals. Mean ± SD, *p<0.05, student’s *t* test. (C) A novel KLRG1^+^ TIGIT^+^T-bet^dim^Eomes^hi^ CD8^+^ T cell population is significant up-regulated in HIV infected individuals. KLRG1 and TIGIT double negative or double positive cells was extracted from B. Expression of T-bet and Eomes was analyzed. The percentage of KLRG1^+^ TIGIT^+^T-bet^dim^Eomes^hi^ was shown. Mean ± SD, *p<0.05, ***p<0.001, ns, not significant, student’s *t* test. (D-F) Blocking KLRG1 restores the activation of T cells to HIV peptides stimuli. *Ex vivo* PBMCs from chronically HIV-infected individuals were stimulated with HIV Gag/Nef peptide pool in the presence of isotype, KLRG1, or PD1 blocking antibodies. (D) Representative flow cytometry plots with gating for CD8 T cells, showing IFN-γ and TNFα responses of PBMCs from an HIV-infected individual. No HIV-1 Gag stimulation with an isotype control is shown as a negative control. A positive control with PMA and ionomycin (1:500) treatment is shown. (E and F). The frequency (%) of IFN-γ (E) and TNFα (F) positive CD8^+^ T cells (n = 8) is shown. *p* values were calculated by Wilcoxon matched-pairs signed ranked test. See also Figure S3

**Figure 4.**
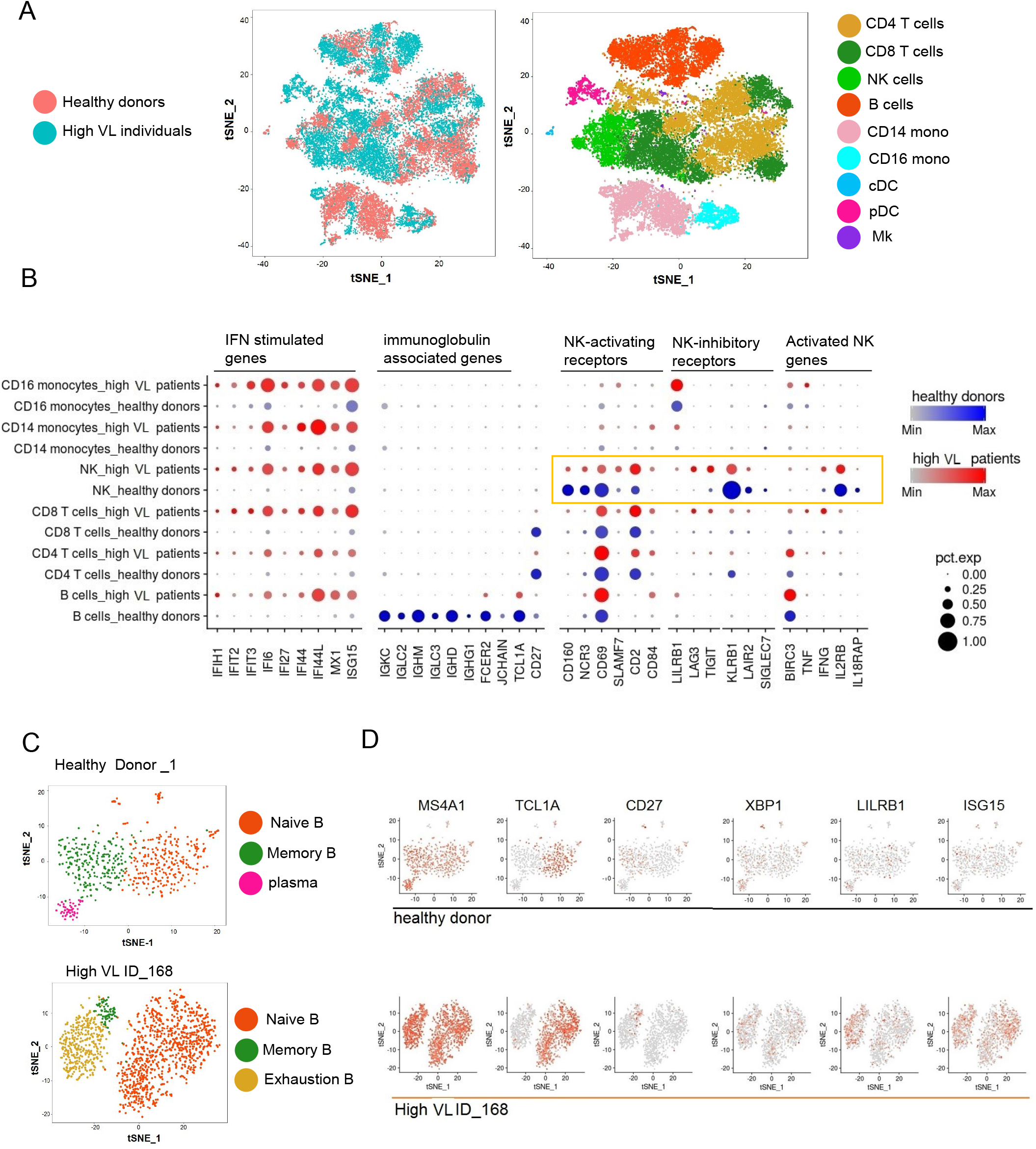
Integrated Analysis of Cell Clusters Identified in PBMCs from Healthy and HIV-Infected Donors. (A) tSNE plots of integrated datasets from the healthy and HIV-infected donors (left) and identification of nine major cell subpopulations (right). NK, natural killer cells; CD14 mono, CD14+ monocytes; CD16 mono: CD16+ monocytes; cDC, conventional dendritic cells; pDC, plasmacytoid dendritic cells; Mk, megakaryocytes. (B) Expression of ‘variable genes’ in six of the cell clusters in healthy and HIV-infected donors. The color intensity indicates the average expression level in a cluster and the circle size reflects the percentage of expressing cells within each cluster. (C) tSNE projections of B cell clusters from healthy donor HD_1 and HL-HIV-infected donor ID_168. (D) Feature plots show the distribution of B cell marker genes in the B cell subsets indicated in (C). Red and gray dots represent cells with high and no/low gene expression, respectively. See also Figure S4

To delineate the relationship between up-regulated KLRG1 and T cell dysfunction, we evaluated whether KLRG1 blockade on HIV-specific CD8^+^ T cells could restore the antiviral immune responses. PBMCs from HIV-infected individuals were stimulated with HIV Gag/Nef peptide pool and treated with KLRG1 blocking or isotype antibodies, and then IFN-γ and TNFα expressing cells were evaluated to determine the restoration of HIV-specific T cell function. In the representative sample, incubation with KLRG1 blocking antibody (20ug/mL) significantly increased the percentage of IFN-γ expressing HIV-specific CD8^+^ T cells from 0.21% to 2.65% (Figure 3D). Meanwhile, the percentage of TNFα expressing HIV-specific CD8^+^ T cells increased from 0.26% to 1.31% (Figure 3D). On the other hand, PD-1 blocking antibody slightly increased the percentages. The statistical analysis for all of the PBMCs isolated from 8 HIV-infected donors showed that KLRG1 blockade could significantly restore cytokine responses (Figure 3E and F).

Previous study described that repetitive and persistent antigen stimulation induced the up-regulation of KLRG1 in virus-specific CD8^+^ T cells ^49^. KLRG1, as an inhibitory receptor, inhibits T cell and NK cell function when activated by its legends ^50,51^. In addition, another inhibitory receptor TIGIT which marks exhausted T cells in HIV infection coexpressed with KLRG1. Also, KLRG1^+^ CD8^+^ T cells showed decreased ability to secret cytokines upon HIV stimulation compared with KLRG1^−^CD8^+^ T cells ^52^. Therefore, it is reasonable that KLRG1 expression was induced by HIV chronic infection to impair the immune system. T-bet and Eomes are T-box transcription factors which regulate the expression of inhibitory receptors and the exhaustion of CD8^+^ T cells^53^. HIV chronic infection down-regulates T-bet and up-regulates Eomes which contributes to poor HIV-specific CD8^+^ T cell functionality ^48^. The significant increase of T-bet^dim^ Eomes^hi^ population in KLRG1^+^TIGIT+ CD8^+^ T cells in HIV”infected PBMCs compared with healthy PBMCs was consistent with these conclusions. Our KLRG1 blockade functional experiments further validate the contribution of KLRG1^+^ population to CD8^+^ T cell exhaustion. Although, KLRG1 was a well”known differentiation maker for lymphocytes, it was not dispensable for T cell and NK cell development and function after virus infection *in vivo* ^54^. It suggests that KLRG1 is a promising and novel therapeutic target for HIV infection.

### HIV Infection Perturbs Immune Cell Composition and Induces B and NK Cell Dysfunction

To extend our investigation to non-T cells, we performed an integrated analysis of the composition of PBMCs from the two healthy donors and three HL-HIV-infected donors. Whereas the cell clusters identified in PBMCs from the healthy donors were similar and overlapping, suggestive of a homogeneous cell composition (Figure S3A), we observed marked cluster heterogeneity in the HIV-infected donors (Figure S3B). This heterogeneity could reflect variations in the individual donor immune response to HIV infection or to different stages of HIV pathogenesis. Interesting, a completely separated tSNE plot with rarely overlapping cells was generated when the healthy donors and the high viral load individuals were compared. Nine clusters were identified with common marker genes in the integrated plots (Figures 3A and S3C). Among the varied genes in the integrated plots, IFN-stimulated genes, such as *IFIH1, IFIT2, IFI6, IFI27, IFI44, IFI44L, MX1 and ISG15*, were significantly up-regulated in monocytes and CD4^+^, CD8^+^, B, and NK cells (Figure 3B). Sustained IFN signaling has been reported to be a key driver of immunosuppression, as demonstrated by CD4^+^ T cell depletion and CD8^+^ T cell expansion during HIV infection ^55,56^. Blocking of the IFNα/β receptor in cART-suppressed HIV-infected humanized mice reversed HIV-induced type-I IFN and rescued the T cell response ^3^. Therefore, suppression of IFN signaling is a potential therapeutic strategy in HIV-infected patients with chronic inflammation.

HIV infection is known to hyperactivate B cells but, paradoxically, it also suppresses the antibody response ^55^. We found that PBMCs from HIV-infected donors contained an increased percentage of B cells compared with healthy donors (22.1%, 26.9%, 33.7% versus 8.7% and 15.1%; Figure S1G). However, compared with healthy donors, expression of immunoglobulin genes, including *IGKC, IGLC2, IGHM, IGLC3, IGHD, IGHG1, FCER2*, and *JCHAIN*, the naïve B cell marker *TCL1A*, and the activated B cell marker *CD27* was abolished or dramatically down-regulated in HIV-infected donors (Figure 3B). In contrast, the B cell inhibitory receptor *LILRB1* was up-regulated in PBMCs from HIV-infected donors (Fig 3B).

In HIV-infected individual ID_168, infection induced a marked expansion of naïve B cells (*TCL1A*^+^ CD27^−^), reduced the memory B cell population (*TCL1A^−^ CD27*^+^), and virtually eliminated plasmablasts (*TCL1A^−^ CD27^−^ XBP1^hi^*) (Figures 3C and 3D). However, the HL-HIV-infected donor PBMC also contained a novel subcluster of B cells characterized by a lack of *TCL1A* or *CD27* and high expression of inhibitory receptor *LILRB1* (also known as CD85j) ^20^, suggesting that this subcluster may represent exhausted B cells (Figures 3C and 3D)^57^. These observations indicate that HIV infection induces B cell dysfunction, possibly secondary to the chronic inflammation and abnormal T cell responses in these patients ^58,59^.

Growing evidence supports a role for NK cells in the antiviral response to HIV-1 ^60^. NK cell cytokine secretion and cytotoxic functions are controlled by multiple activating and inhibitory receptors ^61^. Of note, we found that HIV infection markedly altered the expression of several NK receptor genes (Figure 3B). Among the activating receptor genes, *CD160, NCR3*, and *CD69* were down-regulated while *SLAMF7*, *CD2*, and *CD84* were up-regulated. Conversely, the inhibitory receptor genes *LILRB1, LAG3*, and *TIGIT* were up-regulated and *KLRB1, LAIR2*, and *SIGLEC7* were down-regulated. In addition, we observed increased expression of *BIRC3*, *TNF*, and *IFNG*, which have been proposed to be NK cell activation markers ^22^. Despite the beneficial roles of NK cells in antiviral responses, their function is known to be impaired by chronic HIV infection ^60^. Consistent with this, we found a reduction in the expression of genes involved in cytokine signaling proteins (e.g., *IL12RB* and *IL18RAP)*. Costanzo et al. also observed a reduction in *IL18RAP* expression in NK cells in HIV-infected donors, and this was rescued in individuals inoculated with an HIV vaccine ^22^. These data indicate that chronic HIV infection is associated with an impaired NK cell response.

Collectively, our results underscore the ability of scRNA-seq to overcome obstacles presented by the complexity and heterogeneity of the immune system to enable investigation of pathophysiological changes in very small cell populations ^26^. In this study, we applied scRNA-seq to analyze how HIV infection perturbs the immune cell transcriptome, and we identified marked changes in the composition and function of CD4^+,^ CD8^+^ T, and B cell subsets as well as the function of NK cells. Importantly, exhausted CD8^+^ T cell populations expressing *CD160, TIGIT*, and *KLRG1* were identified here in three HL-HIV-infected individuals. KLRG1 co-expressing with TIGIT marked a new exhausted population and KLRG1 blocking antibody reversed the cytokines response of virus-specific CD8^+^ T cells. This validated the contribution of KLRG1^+^ population to exhaustion and suggested KLRG1 could be a novel target for immunotherapy against HIV infection.

The immune cell exhaustion landscapes demonstrated by this and previous studies ^62^ provide a clearer illustration of HIV-induced immune deficiency. Chronic proinflammatory signaling induced by HIV infection and replication leads to activation of the immune system; however, the accompanying depletion of CD4^+^ T cells reduces their effector functions and, in turn, impairs the CD4^+^ T cell-dependent functions of CD8^+^ T cells and B cells. Thus, despite the fact that CD8^+^ T cell and B cell populations can be activated and expanded during HIV infection, the cells cannot differentiate into effector cells to counter the infection. Moreover, T and B cell exhaustion exacerbates the immune deficiency ^55,56^. Taken together, our data support a vital role for chronic inflammation and immune cell exhaustion in the pathogenesis of HIV infection.

## EXPERIMENTAL PROCEDURES

### Ethical Statement

Written informed consent was obtained from all donors or a parent under a study protocol approved by the Human Research Protection Program at UCSD.

### Human Subjects and PBMC Isolation

Blood from healthy donors and PBMCs from HIV-infected donors were obtained from the UCSD AntiViral Research Center. Blood samples were isolated by Ficoll density centrifugation (Invitrogen) and PBMCs were removed, washed, and frozen until analyzed. scRNA-seq was performed on PBMCs from six HIV-infected donors, three each with high or low viral loads, and one healthy donor. In addition, we also analyzed a published scRNA-seq dataset (from 8381 PBMCs) from another healthy donor (10x Genomics).

### scRNA-seq Library Construction

PBMCs from the seven donors were thawed, and dead cells were removed using magnetic beads (AMSBIO, CB002) to ensure cell viability >80%. scRNA-seq libraries were prepared on the 10X Genomics platform using Chromium™ Single Cell 3’ v2 Reagent Kits and a Chromium instrument according to the manufacturer’s protocol. Libraries were sequenced on a HiSeq2500 platform to obtain 100-bp and 32-bp paired end reads using the following read length: read 1, 26 bp; read 2, 98 bp; and i7 index, 8 bp. The libraries were sequenced to reach ~50,000 reads per cell.

### scRNA-seq Mapping

The scRNA-seq dataset was mapped against the human hg19 reference genome using CellRanger (v2.1; 10x Genomics). GRCh38-1.2.0 was used as a gene model for the alignment and was provided as part of the CellRanger pipeline as a compatible transcriptome reference. The Cell Ranger count function from the CellRanger pipeline was used to generate scgene counts for a single library. This pipeline was also used for alignment, filtering, and UMI counting.

### Single-Cell Differential Gene Expression Analysis

Secondary analysis on the raw counts generated from the Cell Ranger pipeline was performed using Seurat (v.2.2.0), an R package for single-cell genomics. The analysis was based on the pipeline presented in the Guided Clustering Tutorial by the Sajita Lab. First, the cells were filtered to exclude cells with a large number of genes or with a large amount of mitochondrial DNA. The data were then normalized using a global-scaling method known as “LogNormalize,” which normalizes gene expression measurements for each cell by total expression, multiplies the values by a scale factor, and log-transforms the results. The average expression for each gene was calculated to identify genes with highly variable expression and then scaled to remove background variation. Linear dimensional reduction principal component analysis was performed on the identified variable genes. A graph-based clustering approach was used to cluster the cells by type. The resolution parameter was altered for each sample according to cell number. Typically, a resolution >0.8 was used to identify a large number of clusters so that even small cell populations could be identified. Clusters of the same cell type were recombined when the clusters were labeled. Non-linear dimension reduction (tSNE) was used to visualize the data, and the plots were color-coded by cell type.

### Unsupervised Clustering of PBMCs and T Cells by Specific Markers with Seurat

Genes enriched in a specific cluster were identified by the mean expression of each gene across all cells in the cluster. Then the expression of each gene from the cluster was compared to the median expression of the same gene from cells in all other clusters. Genes were ranked based on the expression level, and the most enriched genes from each cluster were selected.

### Identification of Signature Genes in Exhausted T Cells from HIV-Infected Donors

T lymphocyte clusters were selected in each sample using Seurat vignettes. Additional rounds of calculating highly variable genes, scaling the data, and re-clustering the cells were performed on the T cell subsets. Upregulated and down-regulated differentially expressed genes for each cluster were identified. T cell subsets were identified by the expression of cell type-specific markers in feature plots and heatmaps. Exhausted CD8^+^ T cell clusters were also identified using known exhaustion markers. Exhausted CD8^+^ T cell and effector memory CD8^+^ T cells from three donors with high HIV loads were separately analyzed. All pre-processing procedures and clustering were also performed on the distinct subpopulations. Analysis of differentially expressed genes in these two cell types enabled identification of signature genes from the heatmap and feature plots of these subpopulations.

### Pseudotime Analysis

Seurat data matrices were imported into monocle3 for pseudotime analysis ^45,46^. The single-cell dataset was normalize and pre-processed using the DelayedArray packages in Bioconductor and the preprocessCDS function from monocle3, with a num_dim = 10. The dataset then went through one round of dimensionality reduction using UMAP to eliminate noise. The cells in the dataset were partitioned into supergroups for future distinct trajectory recognition. The supergroups were organized into trajectories using SimplePPT. The trajectory plots were visualized and colored by cell types and pseudotime separately, with the start of the pseudotime set to be CD8 T effector memory cells.

### Integrated Analysis of PBMCs from Healthy and HIV-Infected Donors

Using the Seurat integration pipeline ^63^, the PBMC datasets from three donors with high HIV load and two healthy donors were combined by executing the following steps. Individual datasets were filtered, log-normalized, and scaled using the same strategies and parameters as above. After preprocessing the data and selecting genes, canonical correspondence analysis (CCA) analysis was performed to define the shared correlation space between the two healthy donor samples. Multi-CCA was performed on the three samples from high viral load donors. After non-overlapping subpopulations were identified, the CCA subspaces were aligned with dimensional reduction. Another round of clustering was then performed on the two Seurat objects. The two integrations were then combined using the same procedures as above. To detect conserved and differentially expressed genes markers across datasets, differential expression testing was performed on the integrated datasets from all five samples. By performing these two steps, further information was obtained about exhaustion markers, and dot plots were generated to display conserved and differentially expressed gene markers.

### Flow Cytometry

For phenotyping analysis of PBMC, cryopreserved PBMCs were rapidly thawed at 37°C and resuspended in complete RPMI 1640 medium (Life Technologies) supplemented with 10% fetal bovine serum (FBS), 1% penicillin-streptomycin (Hyclone), and 100 U/ml DNase I (StemCell). The cells were washed in complete medium and stained with the viability dye Zombie Aqua (BioLegend), AF700-conjugated anti-CD3 (OKT3), APC-Cy7-anti-CD8a (RPA-T8) from BioLegend and FITC-anti-TIGIT (MBSA43), PE-anti-KLRG1 (13F12F2) (from Thermo Fisher). Cells were washed with staining buffer (BioLegend) and permeabilized by intracellular staining permeabilization wash buffer (BioLegend). Intracellular factors, T-bet and Eomes, were stained by APC-anti-T-bet (4B10) and PE-eFluor 610-Eomes (WD1928) antibody. After all the staining, cells were washed with staining buffer for twice and fixed with FluoroFix buffer (BioLegend), and analyzed on a custom four-laser FACSCanto flow cytometer (BD Biosciences, San Jose, CA, USA). Fluorescence-minus-one (FMO) or isotype staining controls were prepared for gating, and UltraComp beads (Thermo Fisher) were individually stained with each antibody to allow compensation. Data were analyzed using FlowJo version 10 (Tree Star, Ashland, OR, USA).

### KLRG1 Functional Assay

For KLRG1 functional analysis, cryopreserved PBMCs were rapidly thawed and resting in complete RPMI 1640 medium for 12 h. Cells were counted and seeding at a concentration of 1 million/ml. Pooled Gag and Nef peptides (National Institute of Health AIDS Reagent Program) were added into the cells to stimulate HIV-specific cells at a 2.5 ug/ml each peptide. Meanwhile, cells were blocked with isotype IgG (BioXcell), anti-KLRG1 (13F12F2) (Thermo Fisher), or anti-PD1 (Nivolumab) (Selleck) blocking antibody at 20 ug/ml. All of the cells were treated with Brefeldin A to inhibit the secretion of cytokines. After stimulation and blocking for 12 h, the cells were washed twice and then stained for viability with Aqua and CD8a. Followed by permeabilization and intracellular staining of CD3, IFN-γ (EH12.2H7), and TNFα (MAb11) and acquisition on the flow cytometer as above.

## Author Contributions

S.W. and Q.Z. designed and performed the experiments, analyzed the data, and wrote the first manuscript draft; H.H. and K.A. analyzed the data; M.A.Y.K. contributed to obtaining the PBMCs, and data analysis and interpretation; and T.M.R. contributed to the experimental design, data analysis and interpretation, and manuscript writing. All authors contributed to manuscript writing and approved the final version.

## Acknowledgments

We thank Drs. Doug Richman, Mario Stevenson, and Jon Karn for helpful discussions, Kristen Jepsen at the IGM Genomics Center for help with scRNA-seq, Celsa Spina of UCSD CFAR for flow analysis and members of the Rana lab for helpful discussions and advice. We also thank Dr. Song Chen for advice and help in sample preparation and data analysis. This work was supported in part by grants from the National Institutes of Health. Access to retrospectively collected blood samples from HIV-1-infected donors was made possible by the University of California, San Diego Center for AIDS Research, an NIH-funded program (P30 AI036214), which is supported by the following NIH Institutes and Centers: NIAID, NCI, NIMH, NIDA, NICHD, NHLBI, NIA, NIGMS, and NIDDK.

---

**Supplementary Figure 1.**
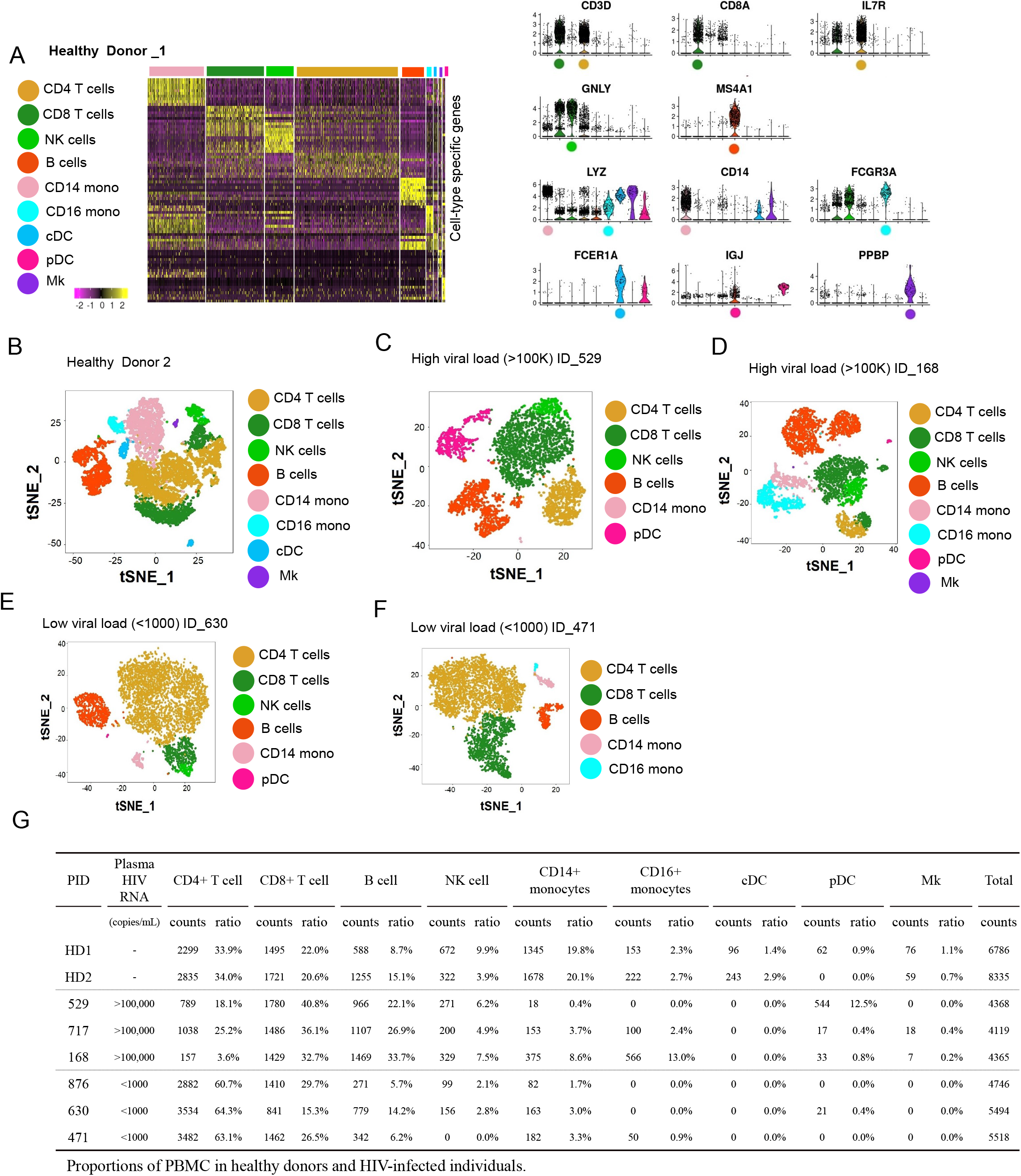
scRNA-seq-Mediated Identification of Cell Clusters in PBMCs from Healthy and HIV-Infected Donors (related to Figure 1). (A) Left: Heatmap of the scRNA-seq dataset from healthy donor_1 showing differentially expressed genes in the indicated cell clusters. Color bar below the map indicates the expression level. Right: Violin plots showing the expression pattern of marker genes for the cell clusters indicated on the left. Each dot represents a single cell and the shapes represent the expression distribution. (B–F) tSNE projections of cell clusters in PBMCs from healthy donor_2 (B), HL-HIV-infected donors ID_529 (C) and ID_168 (D), and LL-HIV-infected donors ID_630 (E) and ID_471 (F). (G) Summary table of the percentage and absolute number each cell type in blood samples from the healthy or HIV-infected donors.

**Supplementary Figure 2.**
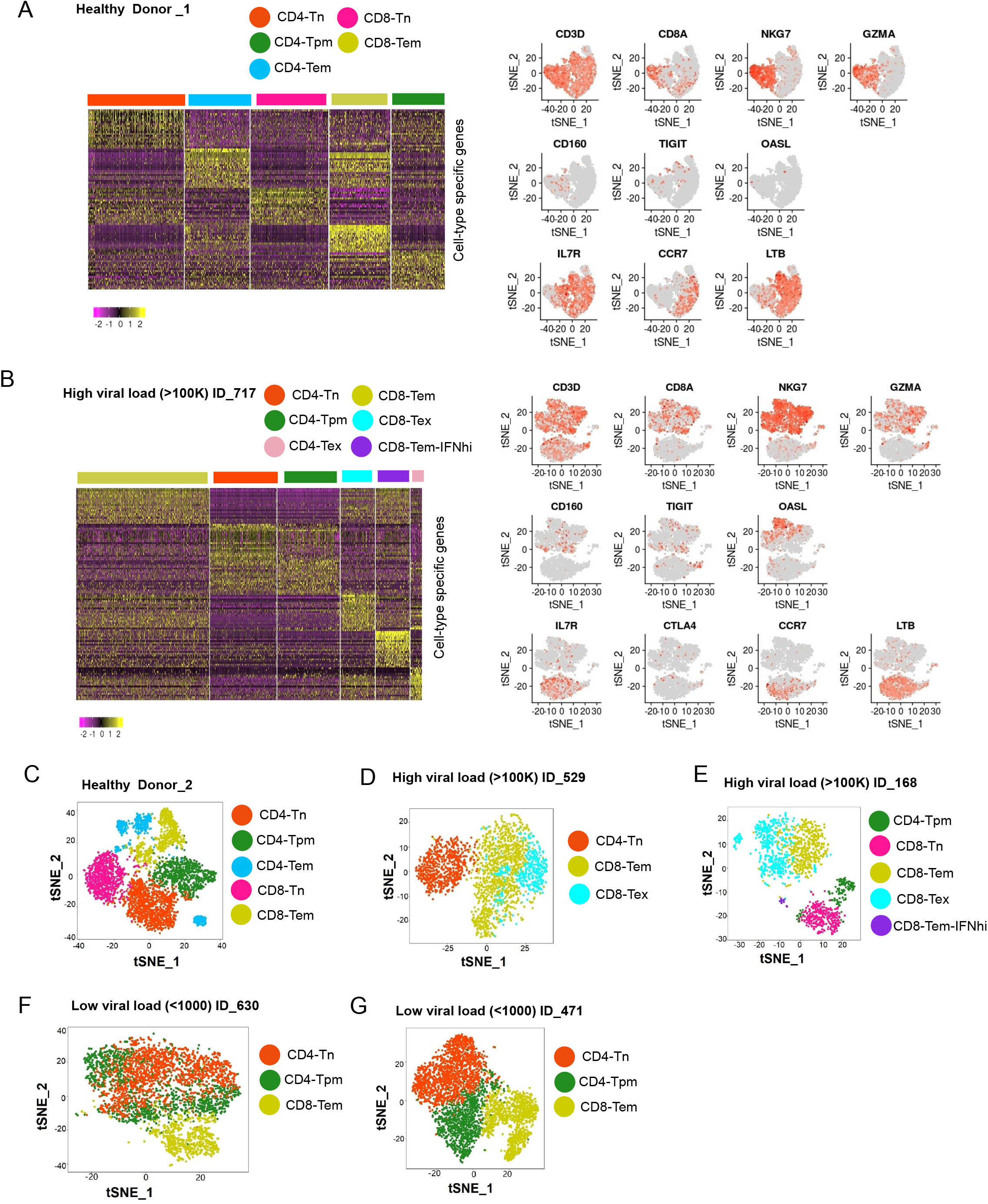
scRNA-seq-Mediated Identification of T Cells in PBMCs from Healthy and HIV-Infected Donors (relates to Figure 1). (A and B) Left: Heatmaps of scRNA-seq datasets showing differentially expressed genes in the indicated T cell clusters from healthy donor_1 (A) and HL-HIV-infected donor ID_717 (B). Color bar below the map indicates the expression level. Right: Feature plots showing the distribution of marker genes for the various cell clusters. Red and gray dots represent cells with high and no/low gene expression, respectively. (C–G) tSNE projection of T cell clusters from healthy donor_2 (C), high-load HIV-infected donors ID_529 (D) and ID_168 (E), and low-load HIV-infected donors ID_630 (F) and ID_471 (G).

**Supplementary Figure 3.**
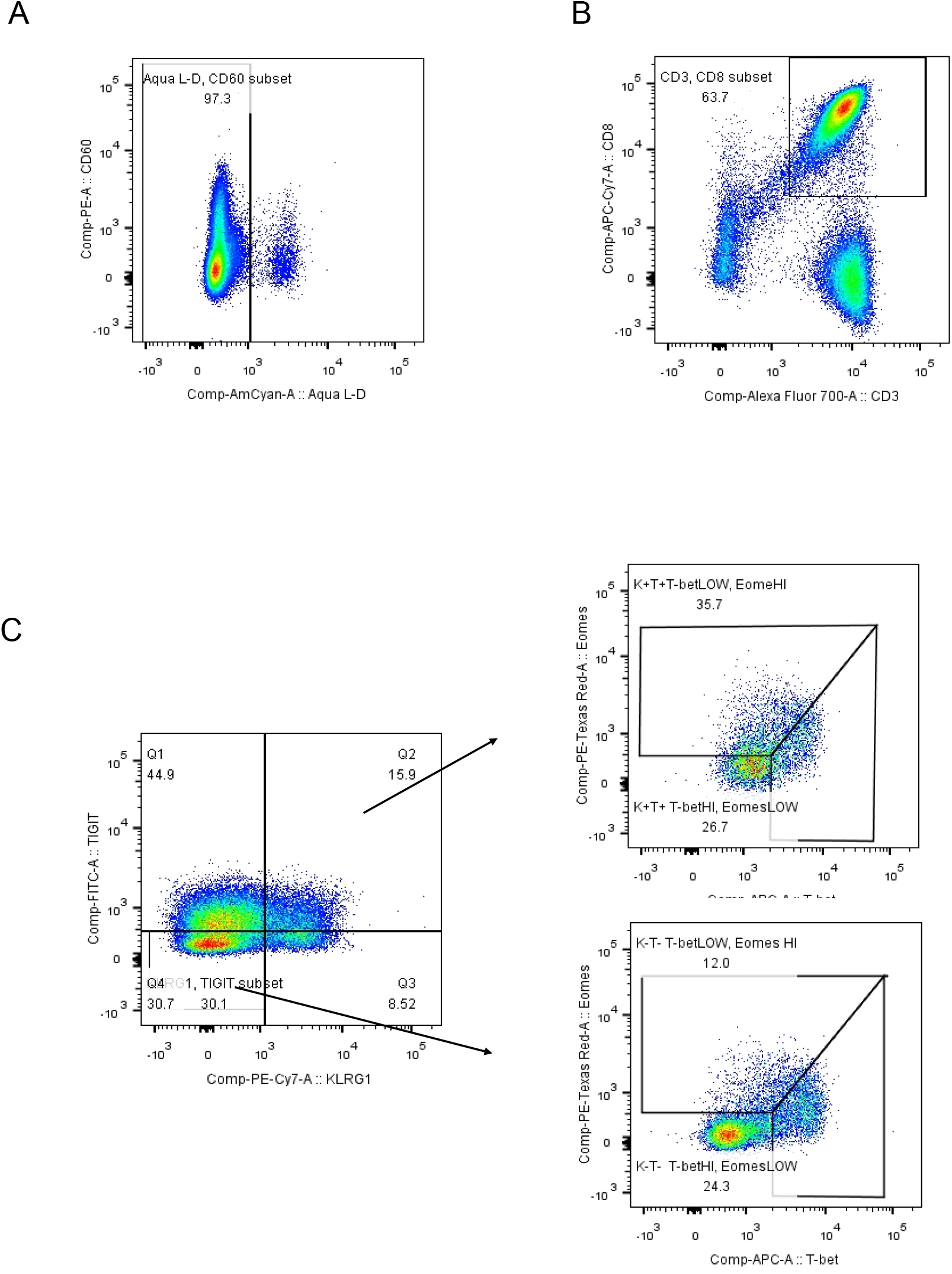
KLRG1 is an exhaustion marker for HIV infected individuals (relates to Figure 3). (A and B) Gating strategy for Aqua negative live cell (A), CD3^+^ CD8^+^ T cells (B). (C) Gating strategy to distinguish the T-bet^dim^ Eomes^hi^ or T-bet^hi^ Eomes^dim^ in KLRG1^+^TIGIT^+^ or KLRG1^−^TIGIT^−^ CD8 T cells.

**Supplementary Figure 4.**
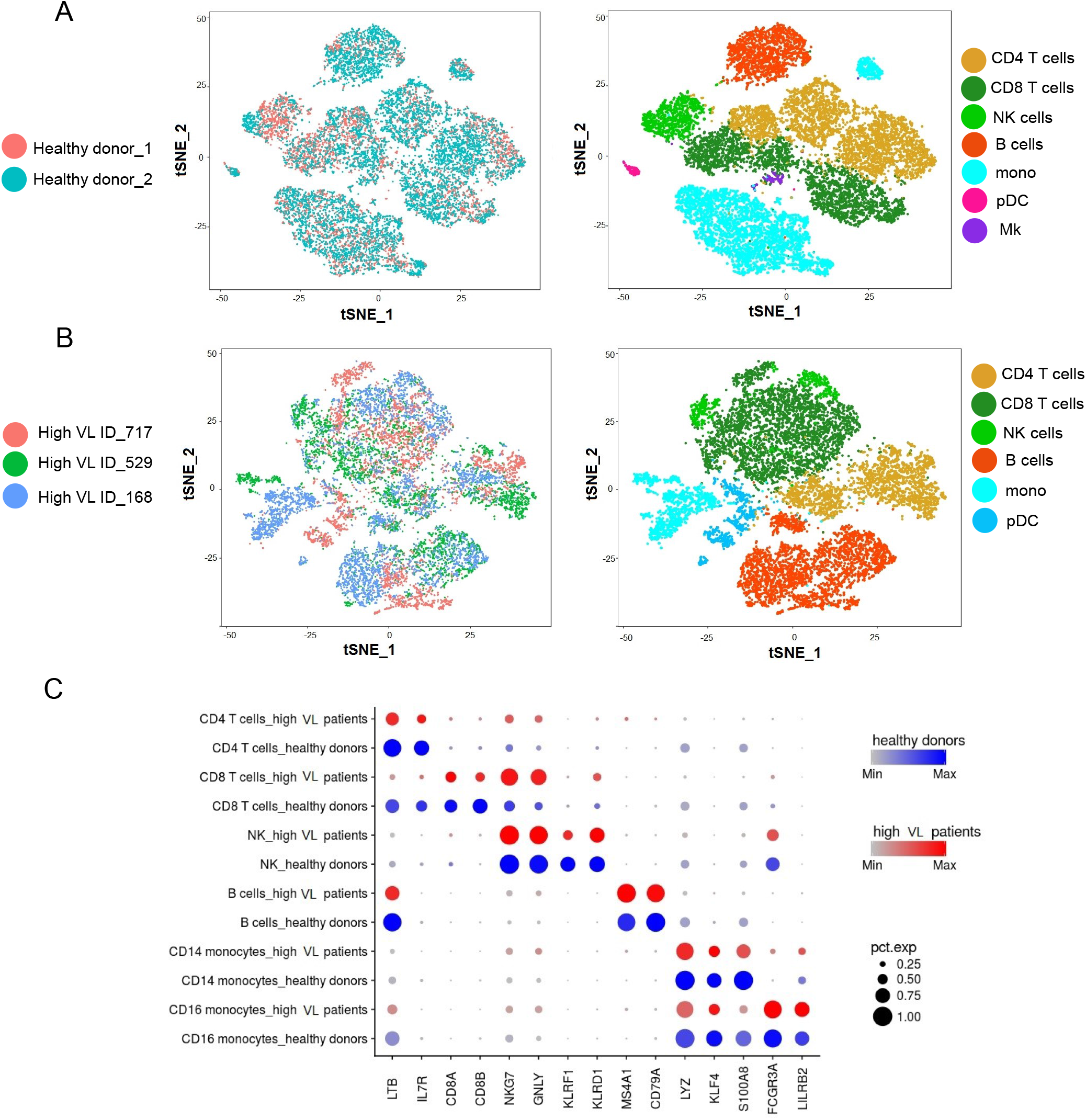
Integrated Analysis of PBMCs from Healthy Donors and High-Load HIV-Infected Donors (relates to Figure 4). (A and B) tSNE plots of integrated datasets from the healthy donors (A) and high-load HIV-infected donors (B). Combined clustering reveals common major subpopulations between samples. (C) Expression of the indicated conserved marker genes in cell clusters from healthy donors and highload HIV-infected donors. The color intensity indicates the average expression level in a cluster and the circle size reflects the percentage of cells expressing that gene within each cluster.

**Supplementary Tables**

**Table S1. Differentially Expressed Genes in the Exhausted T Cell Clusters**

**Table S2. Differentially Expressed Genes in Healthy Donors and High-Load HIV-Infected Donors**

